# Effects of inbreeding and elevated rearing temperatures on strategic sperm production

**DOI:** 10.1101/2024.02.01.578518

**Authors:** Meng-Han Joseph Chung, Md Mahmud-Al-Hasan, Michael D. Jennions, Megan L. Head

## Abstract

Males often strategically modify their rate of sperm production based on the social context, but it remains unclear how environmental and genetic factors shape this plasticity. In freshwater ecosystems, high ambient temperatures often lead to isolated pools of hotter water in which inbreeding occurs. Higher water temperatures and inbreeding can impair fish development, potentially disrupting sperm production. We used guppies (*Poecilia reticulata*) to investigate how developmental temperature (26 °C or 30 °C) and male inbreeding status (inbred, outbred) influence sperm production in the absence or presence of a female (i.e., sperm priming response). We also tested if sperm priming was affected by whether the female was a relative (sister), and whether she was inbred or outbred. A higher rearing temperature had no effect on the rate of sperm production or the priming response. Inbred males produced significantly more sperm in the presence of an unrelated, outbred female than when no female was present. Conversely, outbred males did not alter sperm production in response to female presence or relatedness. In addition, inbred males showed marginally greater sperm production when exposed to an unrelated female that was outbred rather than inbred, but no difference when exposed to an inbred female that was unrelated versus related. Together, only inbred males increased sperm production in response to the presence of a female, but this depended on her being outbred. This suggests stronger sexual selection on inbred males to allocate ejaculate resources, perhaps due to greater benefits when mating with outbred females in better condition.

## Introduction

Males often plastically adjust the rate of sperm production, ejaculate size and/or ejaculate composition based on the social context (Kelly and Jennions 2011; Bartlett et al. 2017). This adaptive plasticity is favored by selection for several reasons. First, sperm production and germline maintenance are energetically expensive (Maklakov and Immler 2016). Evidence for condition-dependence of ejaculate traits (Macartney et al. 2019) suggests that sperm production is costly and trades off with investment in somatic traits (Dowling and Simmons 2012). Second, sperm are vulnerable to oxidative stress due to their high metabolic activity and limited DNA repair (Reinhardt 2007; Helfenstein et al. 2010). This vulnerability results in the deterioration of stored sperm, manifest as slower swimming speed and decreased longevity, which lowers fertilization success (review: Monaghan and Metcalfe 2019). In combination, costly sperm production discourages constant sperm production at the maximum rate, while post-meiotic damage to sperm favors a shorter interval between its production and ejaculation (Pizzari et al. 2008). These factors select for males that adjust sperm production in response to the social context (i.e., sperm priming response; Aspbury and Gabor 2004). Indeed, strategic sperm production in response to mate availability or quality, and to the perceived level of sperm competition occurs in many taxa (review: Magris 2021).

Female availability naturally varies as a result of variation in the environment. Changes in abiotic factors, such as precipitation and temperature, often moderate habitat complexity, resource availability and movement between populations (Ferger et al. 2014; van der Hoek et al. 2022), altering mate encounter rates and the adult sex ratio. A classic example is the approach of spring, coinciding with longer days, higher temperatures and more food, which triggers increased conspecific interactions and reproductive activities (Kahn et al. 2013; De Marchi et al. 2015). However, despite strong correlations between the physical environment and mating opportunities, studies of strategic sperm investment typically manipulate the social context within a single, constant environment (e.g., Moatt et al. 2014; Cattelan and Pilastro 2018; Firman et al. 2018). To date, little attention has been paid to how physical factors interact with the social environment to moderate the plasticity of sperm production.

Temperature is an important environmental factor that can influence the reproductive ecology of species. In ectotherms, temperature can shape mate availability by changing adult sex ratios (Edmands 2021), and through sex differences in behavioral thermoregulation (Ortega et al. 2016). Moreover, warmer temperatures during development can affect the expression of traits that are important for determining resource availability for sperm production, such as body size and metabolism (Baudron et al. 2014). In addition, adult temperature can affect ejaculates and sperm traits (review: Wang and Gunderson 2022). For example, in the European bullhead, adult males living in warmer water show a decline in relative testis size (Dorts et al. 2012); and corals in warmer waters produce fewer sperm (Paxton et al. 2016).

Likewise, males that experience heat stress can exhibit lower sperm quality (e.g., motility, viability) and an increase in the proportion of abnormal sperm (Hurley et al. 2018; Küçük and Aksoy 2020). To date, however, the relative effect of elevated temperatures during development versus adulthood on sperm production is unclear because males in most studies are kept at the same temperature during rearing and adulthood (Zeh et al. 2012; Breckels and Neff 2013; Vasudeva et al. 2014). We therefore know little about how rearing temperature (independent of adult temperatures) affects strategic sperm production in different social environments.

In freshwater ecosystems, warmer temperatures often decrease waterflow and increase the likelihood of isolated bodies of water forming, constraining how conspecifics interact, and elevating the risk of inbreeding (e.g., snail; Jarne et al. 2000). There is ample evidence that inbreeding depresses sperm performance (meta-analysis: Losdat et al. 2014). For example, in zebra finches, inbred males not only sang less often and had less colourful beaks (Bolund et al. 2010), but they also had a significantly lower percentage of normal sperm (Opatová et al. 2016). In addition, there is evidence that stressful environments (e.g., low food availability, high temperatures) exacerbate the detrimental effect of inbreeding on traits related to fitness (Crnokrak and Roff 1999). Despite this evidence, however, we know little about whether inbreeding or stressful environments affect strategic investment in sperm production in response to social cues, with even less knowledge about their potentially synergistic effect.

When living in isolated pools, not only are individuals more likely to be inbred themselves, but the chance of encountering relatives is greater. Animals often exhibit behaviors that reduce the likelihood of mating with kin, such as sex-biased dispersal and mate choice based on kin recognition (review: Nichols 2017). As such, males should invest less in ejaculates when mating with more closely related females to avoid producing inbred offspring with lower fitness (Lewis and Wedell 2009; but see Kokko and Ots 2006 for kin selection benefit). To understand the effect of inbreeding, and its potential interaction with rearing temperature, on strategic sperm production, we designed a study using guppies (*Poecilia reticulata*) to address the following questions: (1) Do males lower their rate of sperm production when in the presence of related females (i.e., inbreeding avoidance)? (2) Do inbred males show less plasticity in sperm production (i.e., inbreeding depression for strategic production in response to social cues)? (3) Do higher rearing temperatures moderate these processes?

The guppy is a freshwater fish in which post-copulatory sexual selection has been widely investigated (Evans and Pilastro 2011). Guppies in tropical streams are often restricted to isolated pools during the dry season (Griffiths and Magurran 1997), leading to higher encounter rates between relatives and naturally elevated levels of inbreeding (Johnson et al. 2010). Inbreeding in guppies has been shown to lower sperm production (Zajitschek and Brooks 2010; Gasparini et al. 2013), fertility (Pitcher et al. 2008; Johnson et al. 2010) and offspring survival (Nakadate et al. 2003). There is some evidence that females actively avoid inbreeding and prefer to mate with unrelated males (Daniel and Rodd 2016), and that multiply mating females bias fertilization toward less closely related males (Gasparini and Pilastro 2011; Fitzpatrick and Evans 2014; but see Evans et al. 2008; Pitcher et al. 2008).

Warm temperatures are common in the tropics where guppies are abundant (Le Roy et al. 2017), but higher temperatures (30°C) can lower sperm performance (Breckels and Neff 2013). A higher sperm count increases fertilization success in guppies (Boschetto et al. 2011), but sperm production is energetically costly (Rahman et al. 2013; Evans et al. 2023).

Consequently, male guppies prudently invest in sperm production in response to female availability (Bozynski and Liley 2003; Cattelan et al. 2016; Cattelan and Pilastro 2018); however, whether stressful environments and/or inbreeding moderate this plasticity is unclear. We examined the interactive effects of male inbreeding status and rearing temperature (26 °C versus 30 °C) on sperm priming response. Furthermore, we manipulated the female’s relatedness (sister or unrelated) and her inbreeding status (inbred or outbred) to test if the risk and/or occurrence of inbreeding affects strategic sperm production.

## Materials and methods

### Fish origin and maintenance

Guppies used in our experiment were descendants of fish from two independent laboratory stocks that both originated from Alligator Creek near Townsville, Australia (19.45°S, 146.97°E). Our stock population has been kept at the Australian National University since 2019. Laboratory-born juveniles were raised in mixed-sex groups until their sex could be determined prior to maturation (an elongated anal fin for males, and a visible gravid spot for females). Males and females were then separated into single-sex 90L tanks (∼50 individuals/tank) to ensure virginity.

All stock fish were maintained under a 14:10 h photoperiod at 26 °C and fed twice daily with *Artemia* nauplii ad libitum and commercial fish flakes. Experimental fish in individual tanks were only fed *Artemia* ad libitum. The project received approval from the Animal Ethics Committee (A2021/04).

### Establishing the inbreeding treatment

To start, we housed a virgin female with a virgin male in a 3L tank. Both individuals were randomly selected from single-sex stock tanks (*n* = 150 pairs). After two weeks the males were removed, leaving the females in the tanks. We inspected tanks daily for newborn fry starting three weeks after the initial pairing. If a female did not give birth within six weeks, or produced fewer than four offspring, she was re-paired with the same male for another week. A total of 80 outbred, full-sibling families were generated. Siblings of the same sex from the same family were housed in communal tanks until they reached maturity. Afterwards, we randomly selected and paired two families to create “blocks” of inbred and outbred fish (e.g., families A and B for block 1, families C and D for block 2; Figure 1). Within each block a male and a female from either the same family (AA or BB) or different families (AB or BA) were paired as above. In total, we generated 60 unique inbred broods and 57 outbred broods spread across 38 blocks.

**Figure 1.**
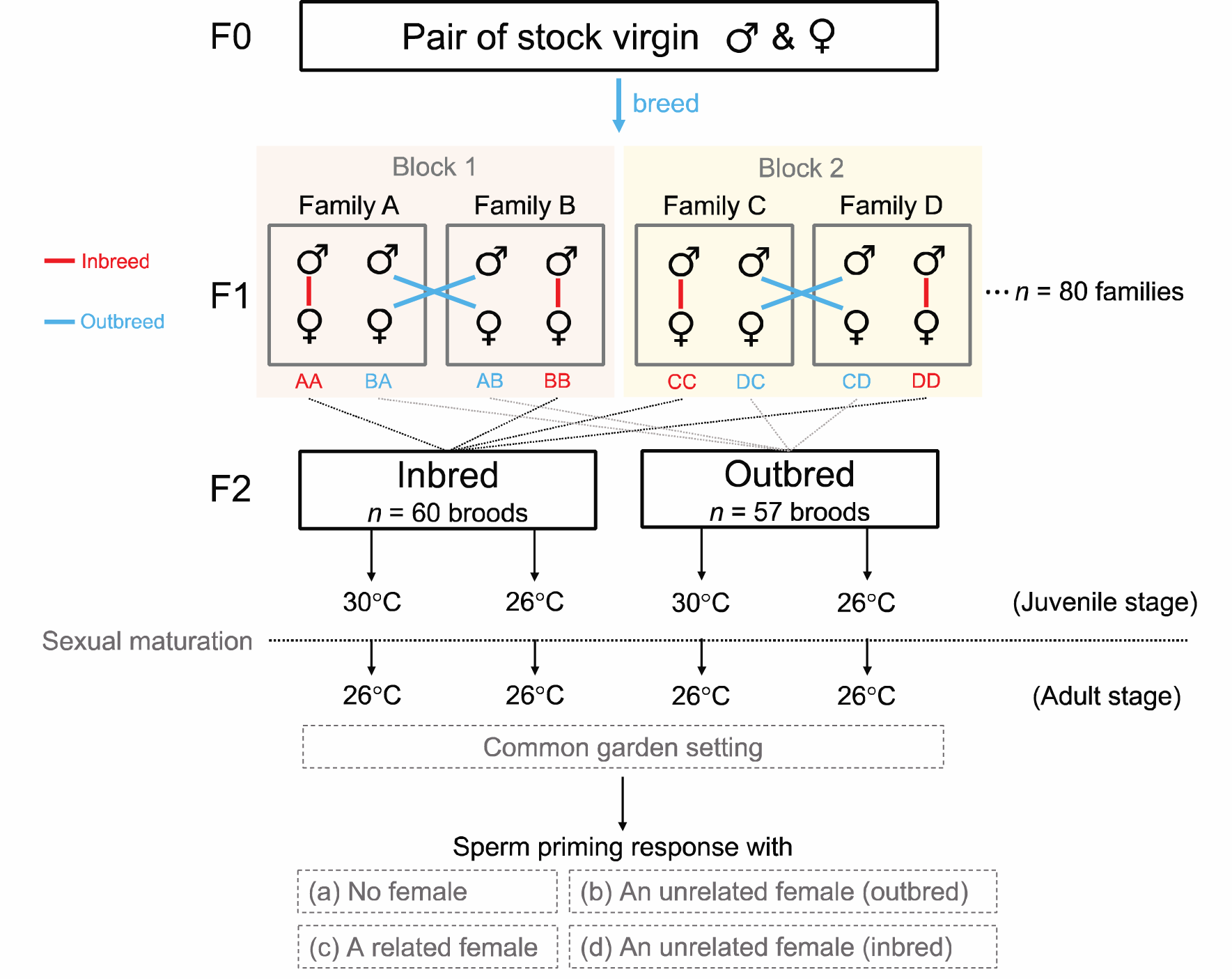
Schematic of the experimental design, showing the block design used to generate inbred and outbred focal fish. Within each block (e.g., Family A × Family B), reciprocal crosses occur between the families (AA, BA, AB, BB). Offspring from each cross-type were assigned alternately to each of the two temperature groups. Two weeks after reaching maturity, adult males were transferred to the common garden temperature and two to five months later exposed to different social contexts to measure their sperm priming response. Note: only inbred males were exposed to environment (d). A related female was either inbred or outbred based on the male’s inbreeding status.

### Manipulating rearing temperature

For each inbred and outbred brood, we randomly assigned half the newborn offspring to either a warm (30°C) or control (26°C) temperature treatment. This split-brood design allowed us to control statistically for variation in genetic and parental effects among broods that might affect trait expression. Offspring were individually housed in 1L tanks, which we checked daily from four weeks after birth onward for fish that had matured. Two weeks after reaching maturation, all fish were placed at 26°C to create a common garden setting for adults. After males had spent two to five months at 26°C, we examined their capacity to modulate the rate of sperm production in response to variation in female availability, female relatedness to the male, and the female’s inbreeding status. This long delay minimised any short-term influence of adult thermal acclimation when the warm-reared (30°C) males were moved to the common garden temperature (Guderley 1990; Seebacher et al. 2014; Little et al. 2021).

### Manipulating female availability and genetic relatedness

To quantify sperm production in response to different social environments, we first emptied a male’s sperm reserves prior to a potential sperm priming phase. Males were anaesthetized with Aqui-S (0.0075% v/v) for 30 s before being placed under a dissecting microscope on a glass slide covered by 1% polyvinyl alcohol solution. The gonopodium (intromittent organ) was swung forwards, and we gently pressed on the male’s abdomen to expel sperm bundles. Following one hour for recovery in their individual 1L tanks, males from each of the four treatment groups (inbred-warm, inbred-control, outbred-warm, outbred-control) (*n* = 85-107 per group; see Supplementary Materials) were randomly introduced into one of three social environments: a 3L tank with either: (a) no female behind a mesh barrier (*n* = 112), (b) an unrelated female (i.e. an outbred, non-sibling) behind a mesh barrier (*n* = 110), or (c) a related female (i.e. his sister, either inbred or outbred based on the male’s own inbreeding status) behind a mesh barrier (*n* = 109) for seven days. We only used stimulus females that were raised at the control temperature to eliminate any temperature-induced variation in female fecundity that might subsequently affect the male’s response.

Inbred males in environment (c) (related female) were unavoidably exposed to an *inbred* sister. Hence, a decreased rate of sperm production could result from either a response to high genetic relatedness to the female and/or a response to her being a lower-quality female (if inbreeding itself lowers female quality; White et al. 2015). To untangle these confounding explanations, we established an additional environment (d) exclusively for inbred males.

Specifically, inbred males experienced the presence of (d) an inbred but unrelated female (i.e., an inbred, non-sibling). Males were assigned alternately to the four environments (a-d) (*n* = 26-31; see Supplementary Materials). Standard length (SL: the snout tip to the base of caudal fin) of the stimulus females ranged from 23.96 to 34.10 mm. Outbred females (27.73 ± 0.13 mm SL) were significantly larger than inbred females (27.15 ± 0.15 mm) (LM, *F*_1,269_ = 7.973, *P* = 0.005). There was, however, no significant size difference between the inbred females that were used in environments (c) and (d) (LM, *F*_1,102_ = 1.141, *P* = 0.288). We were able to control for differences in female inbreeding status and test for an effect of female relatedness on sperm priming for inbred males by comparing how they responded in environments (c) (inbred, related female) and (d) (inbred, unrelated female). For outbred males, the effect of relatedness on sperm priming involved comparing how they responded in environments (b) (outbred, unrelated female) and (c) (outbred, related female).

Males in the no-female treatment did not receive any female cues, while the other males experienced visual and olfactory stimuli from a female. Tanks were separated by white paper to prevent visual contact. Exposing males to a female (or no female) for seven days is a widely used time period in studies of sperm priming in guppies (Bozynski and Liley 2003; Cattelan et al. 2016; Cattelan and Pilastro 2018). It reflects natural conditions as males can be confined to isolated ponds for days to weeks, causing variation in both the availability and genetic relatedness of potential mates (Houde 1997; Magurran 2005). Virgin females show cyclical changes in sexual responsiveness (Liley 1966, 1968), so we mated each stimulus female with a stock male a week before the experiment.

After seven days in the assigned social context, males were anaesthetized and re-stripped to count their sperm (see below). They were also photographed to measure their SL using ImageJ (Abràmoff et al. 2004). We failed to extract sperm from 8 out of 383 males, which was irrelevant to their assigned group (Supplementary Materials). These males were excluded from the analyses.

### Sperm count

The sperm stripped after seven days was collected into a known volume (400-800μL) of saline solution (0.9% NaCl) using a 100-μL pipette. We vortexed the sperm solution for 30 s and mixed it several times using a 20-μL pipette. Next, 3μL of the solution was placed on a 20-micron capillary slide (Leja) to determine the sperm count using CEROS Sperm Tracker (Hamilton Thorne Research, Beverly, MA, USA) under 100× magnification. The samples were collected blind to a male’s treatment (inbreeding status, rearing temperature, social context) and were analysed using the computer-assisted program to eliminate bias. The sperm number was calculated as the mean of five randomly selected subsamples per male (repeatability *r* ± SE = 0.886 ± 0.009, *P* < 0.001, *n* = 375 males).

### Statistical analyses

We ran two separate analyses to address our research questions. First, we ran a linear mixed model (LMM) to investigate the effects of male inbreeding status (inbred, outbred), rearing temperature (warm, control), and social environment (no female, unrelated outbred female, related female) and all three two-way interactions on the rate of sperm production (i.e., number of sperm produced in 7 days).

Second, we noted that inbred males might produce fewer sperm than outbred males in the presence of a related female due to their sister being inbred and therefore of lower quality (e.g., less fecund; White et al. 2015). To test whether the observed effect of male inbreeding status (see *Results*) was confounded by the related female’s inbreeding status, we ran an additional LMM exclusively for inbred males. We separated the effects of inbreeding status of the female and her genetic relatedness to the male by considering three types of female (b,c,d) that inbred males encountered. We compared the response to females that were unrelated and either (b) outbred or (d) inbred to test for an effect of female inbreeding status. We then compared the response to females that were inbred and either (c) related or (d) unrelated to the male to test for an effect of female relatedness. We treated male rearing temperature, female type and their interaction as fixed factors in the model.

In all models, non-significant two-way interactions were removed to test for the main effects of fixed factors (Engqvist 2005). For transparency, all initial and final model outputs are presented in the supplementary materials. We ran Tukey’s post-hoc pairwise test (*emmeans* package) for any significant main effect involving factors with three levels. In all models, brood identity was included as a random factor to account for measurements of several males from the same brood then assigned to different temperatures and social environments. Sperm production is strongly dependent on male size and age (Pitcher and Evans 2001; Gasparini et al. 2010; Kamaszewski et al. 2020), so their SL and adult age at testing were standardized (mean = 0, SD =1) and included as separate covariates in all analyses. Sperm data was power-transformed to fulfill the homogeneity of variances and the normality of residuals assumptions using Levene’s test and Shapiro-Wilks test, respectively.

The significance level was set at alpha = 0.05 (two-tailed). We conducted Wald chi-square tests (*Anova* function in the *car* package) to determine *P* values. Type III sums of squares were used for models with interaction terms, and type II sum of squares for models without interactions. Summary statistics are presented as mean ± SE. Models were run using R v4.0.5 in R studio v1.3.1093.

## Results

Larger males produced significantly more sperm (χ²_1_ = 20.138; *P* < 0.001; supplementary materials Figure S1), while older males produced significantly less sperm (χ²_1_ = 33.986; *P* < 0.001; supplementary materials Figure S2). Controlling for body size and age at testing, sperm count differed significantly between inbred and outbred males depending on the social environment (inbreeding status*social environment: χ²_2_ = 7.195; *P* = 0.027) (Figure 2).

**Figure 2.**
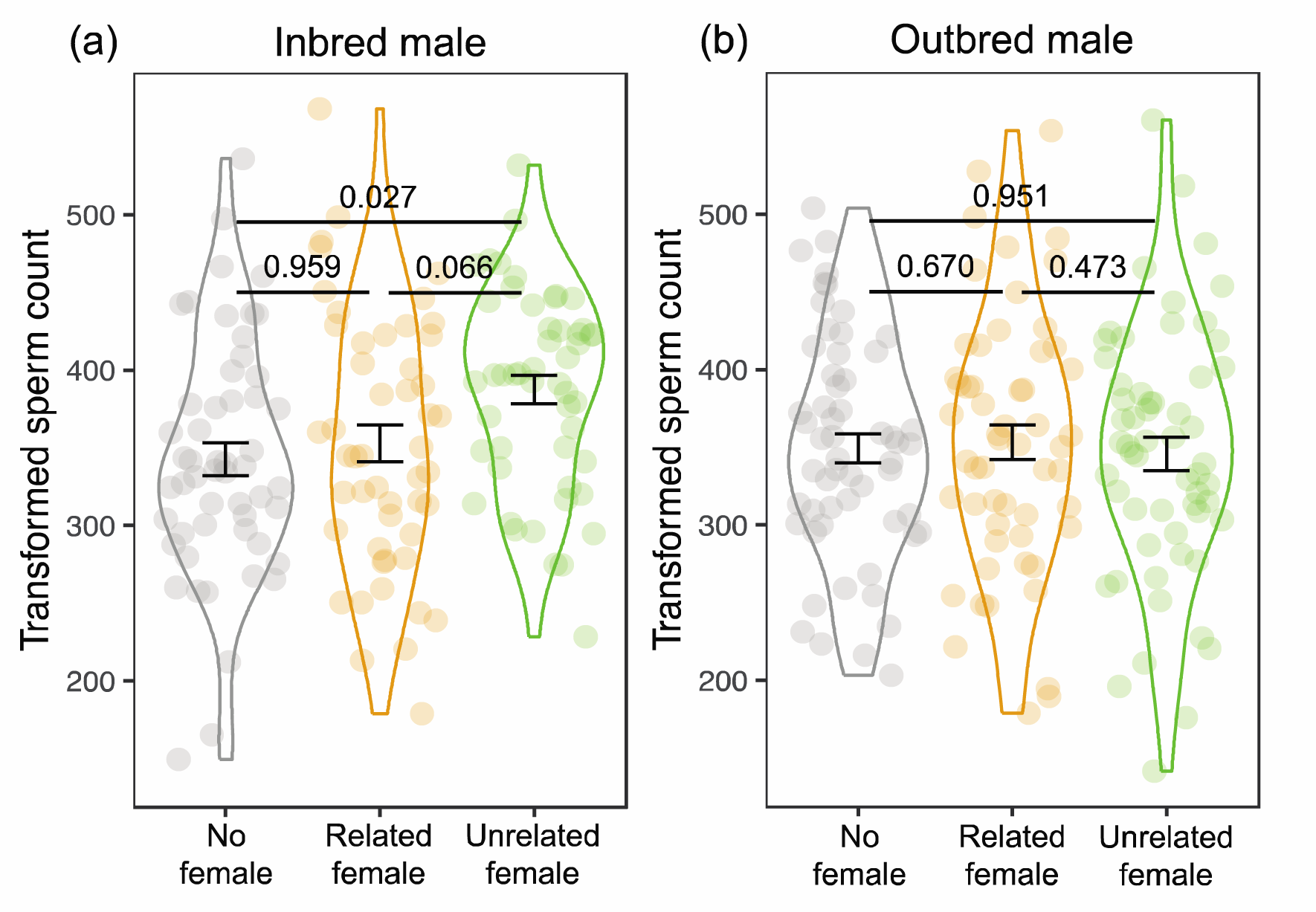
Effect of social environment on sperm count for (a) inbred and (b) outbred males. Grey = no female; orange = related female (inbred or outbred, depending on the male’s inbreeding status); green = unrelated female (outbred). Data is shown as mean ± SE. P values generated using Tukey’s tests are shown.

Neither of the other interactions were significant (inbreeding status*temperature: χ²_1_ = 2.030; *P* = 0.154; temperature*social environment: χ²_2_ = 1.792; *P* = 0.408).

For inbred males, those in the presence of an unrelated female produced significantly more sperm than males without a female (*P* = 0.027). Likewise, inbred males with an unrelated female produced more sperm than those in the presence of a related female (i.e., sister), but this was not significant (*P* = 0.066). The sperm count of inbred males did not differ between the no female or related female treatments (*P* = 0.959) (all Tukey’s tests; Figure 2a). In contrast, the sperm count of outbred males did not significantly differ among the three social environments (Tukey’s tests, all *P* > 0.472) (Figure 2b). These results indicate a sperm priming response by inbred males, when presented with an unrelated female, but not for outbred males. Finally, there was no significant difference in sperm count between males reared at the control temperature and those reared at the warmer temperature (χ²_1_ = 3.062; *P* = 0.080) (Figure 3).

**Figure 3.**
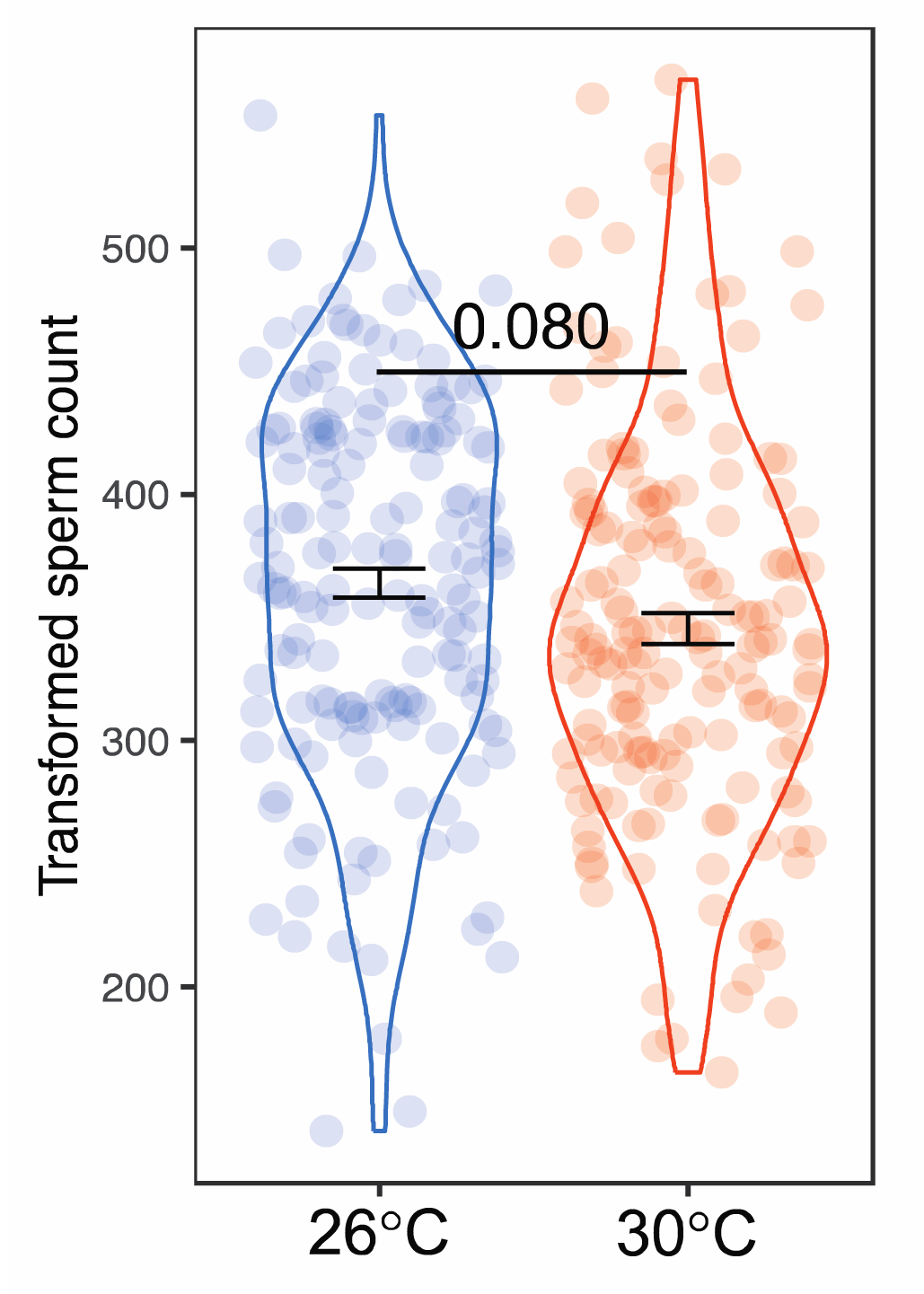
Effect of male rearing temperature on sperm count. Data is shown as mean ± SE. P value shows the significance level of the main effect based on a GLMM (main text).

Given the significant effect of the social environment on the sperm count of inbred males (Figure 2a), we also tested how female relatedness and female inbreeding status affected sperm production by comparing inbred males housed with three types of females (see *Materials and methods*). Controlling for male size (χ²_1_ = 7.573; *P* = 0.006) and age at testing (χ²_1_ = 13.661; *P* < 0.001), female type significantly affected sperm count (χ²_2_ = 6.239; *P* = 0.044), but the effect was not moderated by male rearing temperature (i.e., no interaction: χ²_2_ = 3.795; *P* = 0.150). Despite the significant overall effect of female type, however, we did not find any significant differences in the pairwise comparisons among the three female types (Figure 4). Nevertheless, there was a trend for inbred males exposed to an *outbred*, unrelated female to produce more sperm when compared to inbred males exposed to an *inbred*, unrelated female (Tukey’s tests, *P* = 0.085) (i.e., an effect of female inbreeding status), as well as compared to inbred males exposed to an inbred, related female (Tukey’s tests, *P* = 0.084) (i.e., a combined effect of female inbreeding status and relatedness). In contrast, the amount of sperm produced by inbred males was similar for those experiencing the inbred *unrelated* and inbred *related* female treatments (Tukey’s test, *P* = 0.995) (i.e., no effect of female relatedness). Sperm count for inbred males was unaffected by male rearing temperature (χ²_1_ = 3.264; *P* = 0.071).

**Figure 4.**
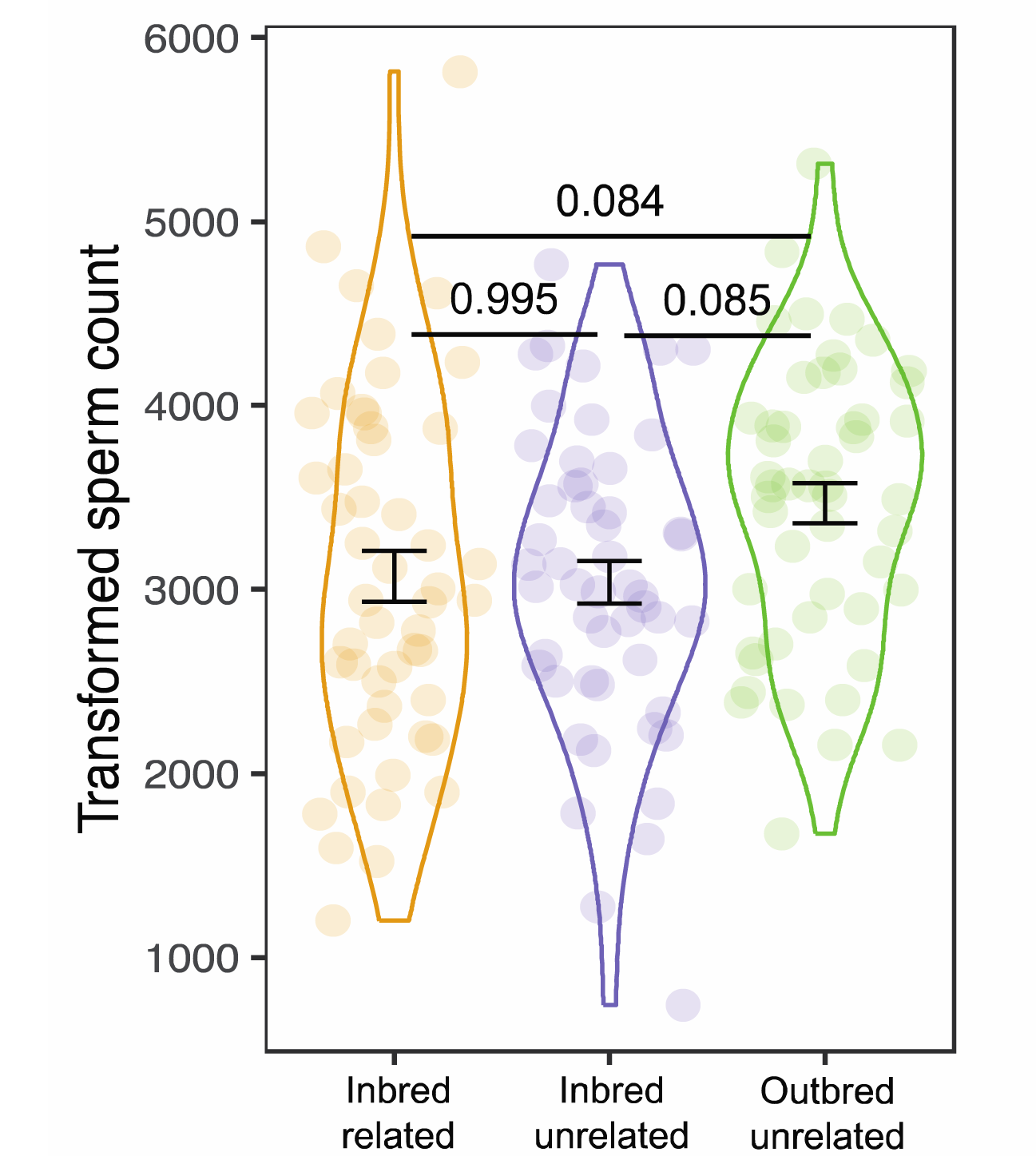
Effect of female type on sperm count by inbred males. Orange = related, inbred female; purple = unrelated, inbred female; green = unrelated, outbred female. Data is shown as mean ± SE. P values generated using Tukey’s tests are shown. Note: there is a significant main effect of female type (main text).

## Discussion

Strategic sperm investment in response to different social contexts has been well studied (Parker and Pizzari 2010; Kelly and Jennions 2011), but few studies have tested how environmental and genetic factors moderate strategic sperm production. This knowledge gap is surprising because variation in the social setting often results from changes in environmental factors (Banks et al. 2005) and inbreeding levels (Kyriazis et al. 2021). Further, synergistic interactions between inbreeding and physical environments (Reed et al., 2012) might amplify the individual effect of each factor on sperm investment, hence fitness. Here, we explicitly disentangled four covarying variables that male guppies in isolated ponds experience, namely – inbreeding, thermal stress, mate availability and mate quality. Male inbreeding status determined how sperm production changed in response to the social context, while a higher rearing temperature did not alter sperm production or sperm priming response.

### Inbred males adjusted sperm production based on female inbreeding status, but outbred males did not

Plasticity in sperm production was only observed for inbred males. Inbred males housed with an unrelated, outbred female produced more sperm than those housed with either no female (*P* = 0.027) or with their sister, who was both related and inbred (P = 0.066). Given that female inbreeding status and relatedness were confounded in this test, we ran a second analysis where we attempted to disentangle the effects of female inbreeding status and her relatedness. This analysis showed no difference in sperm production between inbred males housed with inbred females that were either related or unrelated, but did show that inbred males housed with outbred unrelated females tended to produce more sperm than those housed with inbred females. This suggests that strategic sperm production by inbred males is driven by *female inbreeding status* rather than their relatedness to the female. In a similar study on burying beetles, inbred females preferred to mate with outbred males, while outbred females showed no preference (Pilakouta and Smiseth 2017). In that study, the authors suggest that this pattern could occur if a decline in offspring fitness is greater when inbred males fertilize the eggs of inbred rather than outbred females (Pilakouta and Smiseth 2017), as this would favor inbred females that actively avoid inbred males. In our experiment, inbred female guppies were significantly smaller than outbred females, suggesting that they are less fecund (Reznick and Endler 1982; Auer et al. 2010). As a result, inbred males might benefit by producing more sperm when mating with outbred females as they have more eggs. Our finding for inbred males aligns with past studies in *Drosophila littoralis* where males prefer outbred females and reproductive output was higher for outbred females (Ala-Honkola et al. 2015).

Unlike inbred males, outbred males did not adjust sperm production in response to the social context. As such, female availability did not increase sperm production (i.e., no sperm priming: Evans 2009; but see Bozynski and Liley 2003; Cattelan et al. 2016; Cattelan and Pilastro 2018). Outbred male guppies in our experiment only encountered outbred females (either related or unrelated). Unlike inbred males, however, outbred males did not increase sperm production when in the presence of an outbred female (irrespective of her relatedness). This difference in strategic sperm production between inbred and outbred males might arise if inbred males gain greater marginal benefits than outbred males by upregulating sperm production in the presence of outbred females. For example, inbred males might compensate for their reduced sperm competitiveness when competing for high-quality (i.e., outbred) mates (Zajitschek et al. 2009; Michalczyk et al. 2010).

### Why does female relatedness not affect sperm production by male guppies?

Given lower fertility (Pitcher et al. 2008; Johnson et al. 2010) and reduced offspring fitness (Nakadate et al. 2003) caused by inbreeding in guppies, we expected males to reduce sperm investment when encountering their sister. We offer several possible reasons for the absence of any effect of female relatedness on a male’s sperm production. First, males may use other mechanisms to lower the risk of inbreeding, including male-biased dispersal (Croft et al. 2003; Borges et al. 2022), changes in mating effort (Dougherty et al. 2022) and strategic ejaculation (Wedell et al. 2002; but see Simmons and Thomas 2008). For example, male guppies reduce the intensity of their courtship when directed towards sisters (Fitzpatrick et al. 2014). Second, the males in our study were socially isolated with no interactions with conspecifics prior to testing. Studies of sperm priming often use males reared in mixed-sex groups (Cattelan et al. 2016; Cattelan and Pilastro 2018) or collected from the wild (Aspbury and Gabor 2004; Chung et al. 2019). Prior social experience might be critical to acquire the phenotypic information required for kin discrimination (Penn and Frommen 2010; de Boer et al. 2021). However, this may not be the case for guppies. In an elegant experiment, Daniel and Rodd (2021) showed that male guppies born and reared in isolation could readily discriminate between full and half siblings from different broods. This suggests that early-life exposure to phenotypic cues of kinship is not a prerequisite for kin recognition in guppies.

Third, male guppies may not benefit from post-copulatory mechanisms that reduce inbreeding (Zajitschek et al. 2006; Pitcher et al. 2008) because females show strong mate preferences for unrelated males (Daniel and Rodd 2016); or because there are kin selected benefits to fertilizing sisters, despite inbred offspring being less fit (Kokko and Ots 2006).

Interestingly, instead of exhibiting inbreeding avoidance, there is evidence that male guppies upregulate sperm velocity in the presence of related females (Fitzpatrick et al. 2014), potentially offsetting cryptic female choice against their sperm (Gasparini and Pilastro 2011; Fitzpatrick and Evans 2014). Costly sperm production should discourage inefficient insemination, but mating with sisters can be advantageous when mating opportunities are scarce and there is a low opportunity cost for males (Waser et al. 1986; Kokko and Ots 2006). In this light, it is noteworthy that male guppies in our study and in Fitzpatrick et al. (2014) were virgins, held in sexual isolation prior to testing. This means that the stimulus female was their only apparent mating opportunity, with no alternative options (i.e., no opportunity costs). This might explain why there was no effect of female relatedness on the rate of sperm production. By way of analogy, female guppies biased paternity towards unrelated males only when they received sperm from related and unrelated males simultaneously (Gasparini and Pilastro 2011; Fitzpatrick and Evans 2014). There was no differential usage when females were inseminated by a single related or unrelated male (Gasparini and Pilastro 2011). Our findings raise several questions for further research: Does the mating status of male guppies affect how they respond to females that differ in their relatedness? And what role do opportunity costs play? For example, male guppies discriminate among females that vary in size less pronouncedly when they had previously encountered females consecutively rather than simultaneously (Jordan and Brooks 2012). It would be interesting to test how sperm investment differs for sequential and simultaneous encounters with related and unrelated females (Barrett et al. 2014).

### Warmer rearing temperatures neither affect sperm production nor modify the effect of inbreeding or social context

We found no interaction between male inbreeding status and rearing temperature. This is unexpected as inbreeding depression is exacerbated under stressful environments in many taxa (Armbruster and Reed 2005; Reed et al. 2012). In addition, inbreeding has been proposed as one reason why stocks of captive guppies have a narrower thermal tolerance than wild-caught individuals (Karayucel et al. 2008; Breckels and Neff 2013). Unlike previous studies (Breckels and Neff 2013; Rahman et al. 2020), we found no significant effect of rearing temperature on guppy ejaculates. It is worth noting that males in these earlier studies were maintained at 30°C or even greater temperatures (32°C) during sperm measurement, so their findings may result from effects of the adult rather than developmental environment. We suggest that any negative effect of a high developmental temperature might be reversed after adults are returned to control temperatures for several months. This result is in line with a recent meta-analysis of fishes, which reported greater sensitivity of ejaculates to environmental challenges in adulthood than those during the juvenile stage (Macartney et al. 2019).

Finally, we found no interaction between rearing temperature and social context, suggesting that an elevated developmental temperature does not reduce the ability to plastically adjust sperm production (e.g., the rate of production/transportation of sperm into the testicular duct) (Billard 1986). Given that we found no overall effect of rearing temperature on sperm production, this result is perhaps not surprising. However, it would be interesting to know whether the same results would occur if adults were also tested at 30°C since previous studies imply that adult temperature affects sperm production (Rahman et al. 2020). Ultimately, fertilization success is determined by the overall performance of ejaculate traits under sperm competition (Boschetto et al. 2011; Evans and Pilastro 2011). Future research should therefore examine sperm competitiveness and share of paternity when inbred and outbred males compete, and test for any moderating effect of rearing temperature.

## Conclusion

We found that a male’s inbreeding status affected his plasticity in sperm production. This implies stronger sexual selection on inbred males to strategically allocate ejaculate resources. Further, we show plasticity in sperm production in response to a female’s inbreeding status, rather than whether or not she was a relative (i.e., risk of inbreeding). Together, these results suggest that inbred males seem to face greater demands when it comes to identifying outbred females that might provide greater direct (e.g., more eggs) and/or indirect benefits (e.g., enhanced offspring heterozygosity (Fromhage et al. 2009). This may be due to differences in selection favoring inbreeding-dependent plasticity because the proportion of inbred males and inbred females usually covaries across populations (e.g., inbred males and females are more common in isolated pools than in rivers). Our findings suggest that the negative impact of inbreeding on male fertility, at least sperm quantity, might be less than expected if inbred males are better at fertilizing eggs from outbred females that are more fecund.

## Data availability

Supplementary materials and raw data have been updated as supplementary files for anonymous peer view. They have also been deposited at Mendeley Data (https://data.mendeley.com/datasets/xs3675zk3d/draft?a=d4cce246-d3a5-4a8d-9482-07e826a79b0a) and are publicly available as of the publication date.

## Acknowledgements

We thank the ANU Animal Services staff for help with fish maintenance.

## Ethics

The project received approval from the ANU Animal Ethics Committee (A2021/04).

## Author contribution

All authors conceived and designed the study. MMAH and MHJC conducted the experiment. MHJC analyzed the data and drafted the manuscript, with MDJ and MLH providing critical revisions. All authors approved the final manuscript.

