## Supplementary material for "Effects of inbreeding and elevated rearing temperatures on strategic sperm production"

***Part 1 – Sample size for each manipulated variable***

| Inbreeding status | Rearing temperature | Social environment | *n* |
| --- | --- | --- | --- |
| Outbred males | Warm  (*n* = 85) | No female | 28 |
|  |  | Unrelated female (outbred) | 28 |
|  |  | Related female (outbred) | 29 |
|  | Control  (*n* = 88) | No female | 29 |
|  |  | Unrelated female (outbred) | 31 |
|  |  | Related female (outbred) | 28 |
| Inbred males | Warm  (*n* = 107) | No female | 28 |
|  |  | Unrelated female (outbred) | 27 |
|  |  | Unrelated female (inbred) | 26 |
|  |  | Related female (inbred) | 26 |
|  | Control  (*n* = 103) | No female | 27 |
|  |  | Unrelated female (outbred) | 24 |
|  |  | Unrelated female (inbred) | 26 |
|  |  | Related female (inbred) | 26 |

***Part 2 – Statistical outputs from models with or without non-significant interactions***

**2-1 Likelihood that we extracted sperm from the male**

(a) Initial model including the interaction

|  | | | Estimate | | *SE* | χ²_1_ | | *P* |
| --- | --- | --- | --- | --- | --- | --- | --- | --- |
| Intercept (inbred, control) | | | 9.546 | | 2.515 | 14.401 | | **<0.001** |
| Body length (standardised) | | | -0.036 | | 0.561 | 0.004 | | 0.948 |
| Adult age at testing (standardised) | | | 0.409 | | 0.944 | 0.188 | | 0.665 |
| Male inbreeding status (outbred) | | | 0.415 | | 2.993 | 0.019 | | 0.890 |
| Temperature (warm) | | | -0.147 | | 1.319 | 0.012 | | 0.911 |
| Male inbreeding status (outbred) * Temperature (warm) | | | 21.112 | | 18799.063 | <0.001 | | 0.999 |
| Random effect | Variance | *SD* | | Number of groups | | |  |  |
| Brood ID | 66.270 | 8.141 | | 117 | | |  |  |

(b) Final model excluding the non-significant interaction

|  | | | Estimate | | *SE* | χ²_1_ | | *P* |
| --- | --- | --- | --- | --- | --- | --- | --- | --- |
| Intercept (inbred, control) | | | 9.381 | | 2.567 |  | |  |
| Body length (standardised) | | | -0.130 | | 0.558 | 0.055 | | 0.815 |
| Adult age at testing (standardised) | | | 0.561 | | 0.909 | 0.381 | | 0.537 |
| Male inbreeding status (outbred) | | | 1.186 | | 3.026 | 0.154 | | 0.695 |
| Temperature (warm) | | | 0.690 | | 1.185 | 0.339 | | 0.560 |
| Random effect | Variance | SD | | Number of groups | | |  |  |
| Brood ID | 70.02 | 8.368 | | 117 | | |  |  |

**2-2 Changes in sperm production rates in response to different social environments**

(a) Initial model including all 2-way interactions

|  | | | Estimate | | *SE* | χ² (df) | | *P* |
| --- | --- | --- | --- | --- | --- | --- | --- | --- |
| Intercept (inbred, control, no female) | | | 358.437 | | 12.435 | 830.810 (1) | | **<0.001** |
| Body length (standardised) | | | 18.899 | | 4.176 | 20.481 (1) | | **<0.001** |
| Adult age at testing (standardised) | | | -24.959 | | 4.355 | 32.840 (1) | | **<0.001** |
| Male inbreeding status (outbred) | | | -10.845 | | 15.862 | 0.468 (1) | | 0.494 |
| Temperature (warm) | | | -22.995 | | 14.985 | 2.355 (1) | | 0.125 |
| Social environment (unrelated female) | | | 29.628 | | 15.997 | 3.449 (2) | | 0.178 |
| Social environment (related female) | | | 11.943 | | 16.328 |  | |  |
| Male inbreeding status (outbred) * Temperature (warm) | | | 21.590 | | 15.154 | 2.030 (1) | | 0.154 |
| Male inbreeding status (outbred) * Social environment (unrelated female) | | | -38.153 | | 18.107 | 7.028 (2) | | **0.030** |
| Male inbreeding status (outbred) * Social environment (related female) | | | 6.625 | | 18.636 |  | |  |
| Temperature (warm) * Social environment (unrelated female) | | | 10.132 | | 17.812 | 1.792 (2) | | 0.408 |
| Temperature (warm) * Social environment (related female) | | | -14.209 | | 18.100 |  | |  |
| Random effect | Variance | *SD* | | Number of groups | | |  |  |
| Brood ID | 805 | 28.37 | | 113 | | |  |  |
| Residual | 4137 | 64.32 | |  | | |  |  |

(b) Final model excluding the non-significant interactions

|  | | | Estimate | | *SE* | χ² (df) | | *P* |
| --- | --- | --- | --- | --- | --- | --- | --- | --- |
| Intercept (control, inbred, no female) | | | 354.029 | | 10.815 | 1071.502 (1) | | **<0.001** |
| Body length (standardised) | | | 18.736 | | 4.175 | 20.138 (1) | | **<0.001** |
| Adult age at testing (standardised) | | | -25.563 | | 4.385 | 33.986 (1) | | **<0.001** |
| Temperature (warm) | | | -13.379 | | 7.646 | 3.062 (1) | | 0.080 |
| Male inbreeding status (outbred) | | | -0.674 | | 14.215 | 0.002 (1) | | 0.962 |
| Social environment (unrelated female) | | | 34.363 | | 13.181 | 8.009 (2) | | **0.018** |
| Social environment (related female) | | | 3.709 | | 13.429 |  | |  |
| Male inbreeding status (outbred) * Social environment (unrelated female) | | | -38.072 | | 18.035 | 7.195 (2) | | **0.027** |
| Male inbreeding status (outbred) * Social environment (related female) | | | 7.344 | | 18.610 |  | |  |
| Random effect | Variance | *SD* | | Number of groups | | |  |  |
| Brood ID | 902.8 | 30.05 | | 113 | | |  |  |
| Residual | 4085.6 | 63.92 | |  | | |  |  |

(c) Pairwise comparison for the significant interaction between male inbreeding status and social environment

Inbred males:

| Contrast | Estimate | *SE* | df | *t* ratio | *P* |
| --- | --- | --- | --- | --- | --- |
| No female – Unrelated female | -34.360 | 13.200 | 280 | -2.600 | **0.027** |
| No female – Related female | -3.710 | 13.500 | 303 | -0.275 | 0.959 |
| Unrelated female – Related female | 30.650 | 13.700 | 299 | 2.245 | 0.066 |

Outbred males:

| Contrast | Estimate | *SE* | df | *t* ratio | *P* |
| --- | --- | --- | --- | --- | --- |
| No female – Unrelated female | 3.710 | 12.300 | 257 | 0.302 | 0.951 |
| No female – Related female | -11.050 | 12.900 | 294 | -0.854 | 0.670 |
| Unrelated female – Related female | -14.760 | 12.600 | 278 | -1.169 | 0.473 |

**2-3 Changes in sperm production rate of inbred males in response to different female inbreeding status and female relatedness**

(a) Initial model including the interaction

|  | | | Estimate | | *SE* | χ² (df) | | *P* |
| --- | --- | --- | --- | --- | --- | --- | --- | --- |
| Intercept (control, inbred unrelated) | | | 3105.040 | | 158.510 | 383.722 (1) | | **<0.001** |
| Body length (standardised) | | | 205.430 | | 69.480 | 8.742 (1) | | **0.003** |
| Adult age at testing (standardised) | | | -261.740 | | 71.170 | 13.528 (1) | | **<0.001** |
| Temperature (warm) | | | 32.630 | | 213.630 | 0.023 (1) | | 0.879 |
| Female type (inbred related) | | | 279.200 | | 213.310 | 4.061 (2) | | 0.131 |
| Female type (outbred unrelated) | | | 406.130 | | 208.210 |  | |  |
| Temperature (warm) * Female type (inbred related) | | | -562.380 | | 291.490 | 3.795 (2) | | 0.150 |
| Temperature (warm) * Female type (outbred unrelated) | | | -194.960 | | 291.760 |  | |  |
| Random effect | Variance | *SD* | | Number of groups | | |  |  |
| Brood ID | 143114 | 378.3 | | 57 | | |  |  |
| Residual | 492561 | 701.8 | |  | | |  |  |

(b) Final model excluding the non-significant interaction

|  | | | Estimate | | *SE* | χ² (df) | | *P* |
| --- | --- | --- | --- | --- | --- | --- | --- | --- |
| Intercept (control, inbred unrelated) | | | 3237.690 | | 133.360 |  | |  |
| Body length (standardised) | | | 189.960 | | 69.030 | 7.573 (1) | | **0.006** |
| Adult age at testing (standardised) | | | -262.400 | | 70.990 | 13.661 (1) | | **<0.001** |
| Temperature (warm) | | | -232.640 | | 128.770 | 3.264 (1) | | 0.071 |
| Female type (inbred related) | | | -14.000 | | 151.190 | 6.239 (2) | | **0.044** |
| Female type (outbred unrelated) | | | 318.220 | | 147.700 |  | |  |
| Random effect | Variance | *SD* | | Number of groups | | |  |  |
| Brood ID | 136300 | 369.2 | | 57 | | |  |  |
| Residual | 503812 | 709.8 | |  | | |  |  |

(c) Pairwise comparison for the significant effect of female type

| Contrast | Estimate | *SE* | df | *t* ratio | *P* |
| --- | --- | --- | --- | --- | --- |
| Inbred unrelated – Inbred related | 14 | 152 | 135 | 0.092 | 0.995 |
| Inbred unrelated – Outbred unrelated | -318 | 148 | 117 | -2.146 | 0.085 |
| Inbred related – Outbred unrelated | -332 | 155 | 135 | -2.148 | 0.084 |

***Part 3 – Effect of male body size and age at testing on sperm count***

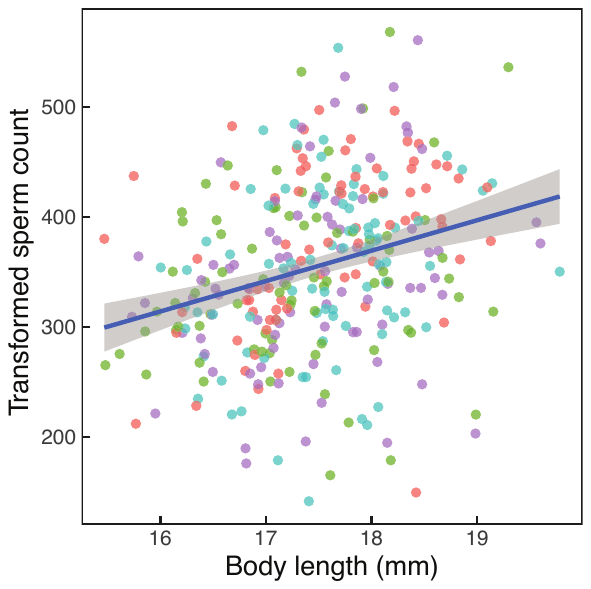

Figure S1. Relationship between male body length and sperm count. Colour indicates the treatment combination of male inbreeding status and rearing temperature: red = inbred-control; green = inbred-warm; blue = outbred-control; purple = outbred-warm. Body size-dependence of sperm count across all males is shown using a regression line with the 95% confidence interval.

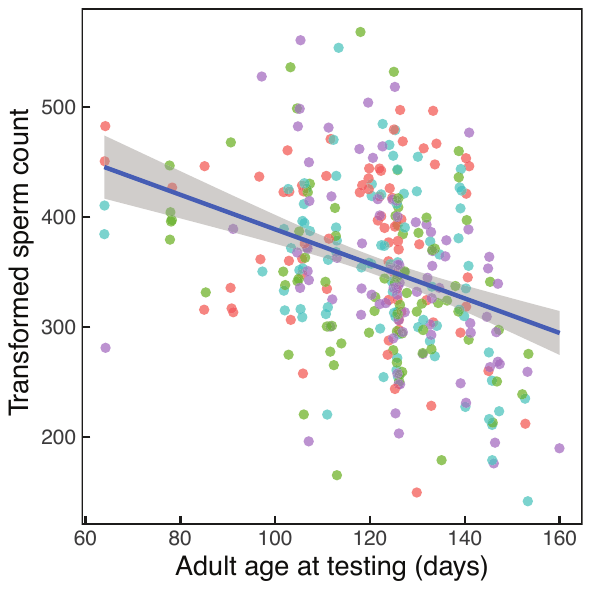

Figure S2. Relationship between male age at testing and sperm count. Colour indicates the inbreeding status-rearing temperature combination of male treatments: red = inbred-control; green = inbred-warm; blue = outbred-control; purple = outbred-warm. A regression line with the 95% confidence interval for all datapoints was shown.
